# From Substance to Computation: Internalization and Hierarchical Structure in Evolutionary Phonology

**DOI:** 10.1101/2020.04.09.033340

**Authors:** Shin-ichi Tanaka

## Abstract

In the history of phonological theory, the paradigm of computational systems has shifted in tandem with a better understanding of substantive issues such as typology, acquisition, social variation, and historical change in sound structure. The paradigm shift in the past 50 years can simply be characterized as the one from ‘serial derivation by rules’ to ‘parallel evaluation by constraints,’ and now Optimality Theory (OT) focusses on substantive issues by improving its phonetic groundings and has ceased groping for a better mode of computation. This is because OT is primarily a substantive theory of CON, and at least in its standard version, the computational systems of Eval and GEN are merely ‘given assumptions.’

In this article, we will overview some arguments against OT in substantive respects. As an ultimate problem in substance, it cannot solve ‘the poverty paradox,’ which means the paradox of ‘the poverty of the stimulus’ in ontogeny and ‘the poverty of the inheritance’ in phylogeny. Moreover, as a proximate problem in substance, certain gaps to be missed in syllable typology would erroneously be predicted to exist by OT. Alternatively, we will rethink the mode of computation in phonology and propose a new paradigm for phonology from the viewpoint of language evolution under a minimalist lens. Our proposal is based on Fujita’s (2016a, 2017) Hypothesis on Motor Control Origin of Merge in Language Evolution, which solves ‘the poverty paradox’ and thus satisfies both explanatory and evolutionary adequacy. We will demonstrate that substantive findings in OT can successfully be carried over to this scenario, in which the empirical problem concerning the typological gap is offered a reasonable explanation. We will also show that phonology has a vital role in computation and is not merely a subsidiary issue at the interface of the Sensory-Motor systems in linearization or externalization. We will take up one case for this claim: the English syllable CCVC has structural ambiguity, which means that phonology involves internalization with some mechanisms in order to create different hierarchical structures.

## 1 Introduction

This article will be devoted to rethinking the role of computation in phonology and elaborating the role of phonology in the strong minimalist thesis (SMT; Chomsky 2004, 2005) from the viewpoint of the evolution and origins of language. While current trends in phonological theory mainly focus on the function of substance in sound structure and have almost ceased examining the nature of computation in phonology, we will claim that phonological theory never satisfies evolutionary adequacy until it seriously re-examines its computation and re-creates a new paradigm for phonology under the minimalist lens.

In the history of phonological theory, the paradigm of computational (formal) systems has shifted in tandem with a better understanding of substantive (empirical) issues such as typology, acquisition, social variation, and historical change in sound structure, since the adequacy of computation is empirically tested by ultimate and proximate issues in substance.^1^ In particular, the paradigm shift in the past 50 years of phonological theory can simply be characterized as the one from ‘serial derivation by rules’ to ‘parallel evaluation by constraints,’ and now Optimality Theory (OT; Prince and Smolensky 1993/2004) focusses on substantive issues by improving its phonetic bases and has almost ceased groping for a better mode of computation. This is because OT is primarily a substantive theory of CON(straint), a universal set of phonetically-grounded constraints on outputs (markedness constraints) and correspondence constraints on input-output or output-output relations (faithfulness constraints). Hence, at least in its standard version, the computational systems of Eval(uation) and GEN(erator) are merely ‘given assumptions’ and left behind.^2^

In this article, we will rethink the mode of computation in phonology and propose a new paradigm of phonology from the viewpoint of language evolution under a minimalist lens (sections 2.1 and 2.2). By comparing our proposed model with OT, we will demonstrate that OT has crucial problems in biological evolution (macroevolution or language origins) and cultural evolution (microevolution or historical change and typology) and so is not evolutionarily adequate. As an ultimate or biological problem in substance, OT cannot solve ‘the poverty paradox,’ which means the paradox of ‘the poverty of the stimulus’ in ontogeny and ‘the poverty of the inheritance’ in phylogeny over the content of UG (section 2.3). This is because it assumes too rich content of UG in the form of CON, EVAL, and GEN, and hence lacks evolutionary adequacy. To take up CON, for example, there are so many, arguably innumerable, constraints assumed in UG, and it is not at all unknown where such varieties of uniquely-human traits come from. On the other hand, as a proximate or historical problem in substance, certain gaps to be missed in syllable typology would erroneously be predicted to exist by OT (sections 3.1 and 3.2). This problem concerns the typology of superheavy syllables in which CVCC implies the existence of CVVC but not vice versa, and yet OT cannot account for this irreversible implicational relation.

Instead, we will propose a new paradigm for phonology, based on Fujita’s (2016a, 2017) Merge-Only Hypothesis of Human Language Evolution, which solves ‘the poverty paradox’ and thus satisfies both explanatory and evolutionary adequacy (sections 2.1 and 2.2). In our model, ‘the poverty of the stimulus’ and ‘the poverty of the inheritance’ will be accounted for by ‘the richness of the Third Factor’ and ‘the Merge-only content of UG,’ respectively, according to Chomskian SMT and Darwinian gradualism. Then, we will demonstrate that substantive findings about typology, acquisition, social variation, and historical change in OT can successfully be carried over to this scenario, in which even the empirical problem concerning the typological gap is offered a reasonable explanation (sections 3.1-3.3). Specifically, we will show an evolutionarily-adequate account that covers empirically more valid patterns of syllable typology than OT and yet carries over other substantive findings from OT. This extensive account will prove to be possible by assuming the three gradual modes of Merge (i.e., pair-Merge, pot-Merge, and sub-Merge) in parallel with Greenfield’s (1991) action grammar, as well as the on/off settings of the interface conditions in the Sensory-Motor systems.

Finally, we will also show that phonology has a vital role in computation and is not merely a subsidiary issue at the interface of the Sensory-Motor systems in linearization or externalization (section 4). We will take up one case for this claim in which a certain effect of identity avoidance crucially depends on the hierarchical structure in the domain of a syllable; namely, in English, the CCVC structure generally has the identity avoidance effect between the second onset and the coda, while the same sequence does not have it on certain conditions, whose difference is accounted for by separate internal organizations. This means that phonology involves internalization with some mechanisms in order to create different hierarchical structures.

## 2 Phonology Under the Minimalist Lens

### 2.1 The Emergence of Human Language Is Not a Miracle

Since the advent of the SMT, the grammar of a particular language has been considered as recursive Merge, in conjunction with the interfaces of the Sensory-Motor (SM) systems for externalization or phonetic interpretation and of the Conceptual-Intentional (CI) systems for internalization or semantic interpretation. Based on the SMT and Fujita’s (2016a, 2017) Merge-Only Hypothesis of Human Language Evolution, Tanaka (2017, 2018a,b) reconstructs a general scenario in (1) for the origins and evolution of language, showing how species-specific human language (UG) emerges from its precursor in macroevolution and how a particular grammar emerges from UG in microevolution.

(1) The Third Factor in Macroevolution and Microevolution

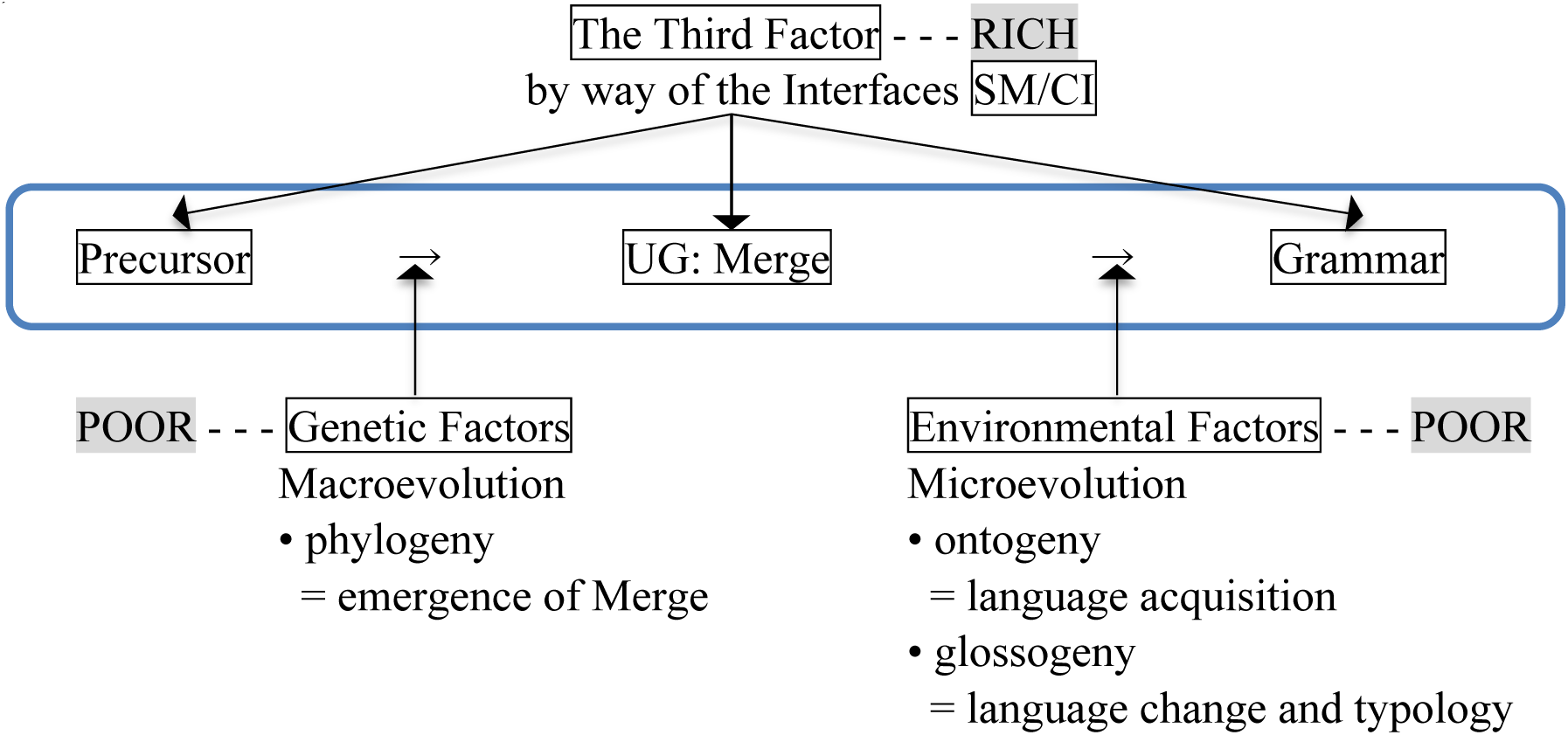

In this model, the phylogeny of human language, i.e., macroevolution, is characterized as the emergence of Merge, which was made possible by some slight mutation in genetics, or what Chomsky (2005) calls “a rewiring of the brain.” It is true that human language emerged from some proto-language by way of Merge, but Merge itself emerged from action grammar as its preadapted precursor by way of parallel evolution, which will be discussed in section 3.1. This is the origins of human language. In microevolution, the ontogeny and glossogeny of a particular grammar emerge from Merge at the SM/CI interfaces and are controlled by environmental and social factors.

What is crucial in (1) is that “the Third Factor” in the sense of Chomsky (2005) inclusively enters into the shift from Merge to the grammar of a particular language as well as the one from the precursor to Merge. The Third Factor is a set of principles of the natural or physical law by which some traits (behaviors or organs) in animals are necessarily governed. In humans, this made the emergence of Merge in macroevolution the way it was. Thus the Third Factor must be the principles in the SM systems exactly because the traits of animals (especially their behaviors) are formed through the SM system. As a consequence, we can have indirect access to such hard issues in macroevolution (i.e., phylogeny) by investigating the grammars of individual languages directly as issues in microevolution.

Another important point is that we do not think of the emergence of human language as a miracle or a skyhook, which appeared out of nothing. Instead, we follow the Merge-Only Hypothesis of Human Language Evolution in assuming that it is within the scope of gradualism although human language has part of a species-specific property. This is because the uniquely-human trait is only Merge, which evolved gradually in parallel with action grammar from neighboring species. ^3^ In other words, a minimal content of UG is a natural feature by which gradualism from other species to humans is maintained.

### 2.2 Solution to ‘the Poverty Paradox’

Chomsky (1986) once raised and answered a significant ontogenetical question concerning human language, known as the logical problem of language acquisition or Plato’s problem: why can human beings acquire a language so early and uniformly, despite very limited learning environments where primary linguistic data are poor in quality and quantity? This is because human beings are endowed with *a priori* knowledge according to the Platonian thesis, meaning the rich content of UG, while environmental factors are poor, meaning ‘the poverty of the stimulus.’

On the other hand, Fujita (2007, 2009) first raised and answered a significant phylogenetical question concerning human language, known as the logical problem of language evolution or Darwin’s problem: why could human beings have evolved a species-specific language, despite very limited genetic environments where uniquely-human linguistic genes are poor in quality and quantity? This is because *a priori* knowledge, Merge, is minimal according to the Darwinian thesis, meaning the poor content of UG, but it has infinite creative capacity although genetic factors are poor, meaning ‘the poverty of the inheritance.’

So ‘the poverty of the stimulus’ implies the rich content of UG, while ‘the poverty of the inheritance’ implies the poor content of UG. So the requirements for language acquisition and language evolution seem contradictory, and linguistic theory with both explanatory and evolutionary adequacy seems impossible. This is ‘the poverty paradox.’

For the Chomskian thesis, environmental factors must be poor, and for the Darwinian thesis, genetic factors must be poor. And we are assuming that UG must be minimal and contains Merge only. So there must be something rich enough to make up for the minimal content of UG, confirm early and uniform acquisition, and lead to the solution of ‘the poverty paradox.’ We argue that this is the richness of the Third Factor by way of the SM/CI interfaces, as given in (1). Human beings share many cognitive capacities within the SM/CI systems with other species which can communicate (Samuels 2011), and these systems are susceptible to the rich principles of the natural and physical law. The only species-specific and domain-specific capacity in humans is Merge.

In short, the Third factor, which works equally in phylogeny and ontogeny, must be rich precisely because it is species-general and domain-general. And if environmental factors are poor, genetic factors in UG are also poor, but the principles of the Third Factor are rich, then we can offer a reasonable explanation to ‘the poverty paradox’ of phylogeny and ontogeny.

### 2.3 Substance-Free or Substance-Bound Phonology

As mentioned in section 1, OT is totally a theory of substance in grammar, because computation in EVAL and GEN have become ‘given assumptions,’ and thus phonologists have focused on CON as a target of inquiry. This means that OT ignores further theoretical development in computation and that phonology goes further toward phonetic experiments and statistic quantifications to test the content or ranking of constraints, an E-language matter. In this sense, OT is totally a ‘substance-bound’ phonology.

However, as is clear from section 2.2, OT suffers from an ultimate or biological problem in substance as it cannot solve ‘the poverty paradox.’ Specifically, it assumes too rich content of UG in the functions of CON, EVAL, and GEN, and hence lacks evolutionary adequacy. To take up CON, for example, there are so many, arguably innumerable, constraints assumed in UG, and it is not at all unknown where such varieties of uniquely-human traits come from. That is, it indeed explains ‘the poverty of the stimulus’ problem in ontogeny but cannot account for ‘the poverty of the inheritance’ issue in phylogeny, which stems from its too rich content of UG.

Alternatively, as we will discuss in section 3.3, substance should be the Third Factor issues by way of the SM systems at the interface, which are beyond the scope of grammar, and of phonological theory. If this is the case, ‘the poverty paradox’ is naturally given a reasonable account. The core issues of linguistic theory are computation, an I-language matter. So the role of phonology is to focus on, and explicitly elucidate, how computation goes in phonological grammar. In this regard, phonology should be “substance-free” in the sense of Hale and Reiss (2000, 2008) and Reiss (2017). This is a logical consequence of language acquisition and language evolution, and let us call such a substance-free version minimalist phonology.

In fact, this minimalist view arguably not only offers phonology with explanatory and evolutionary adequacy, but also even with an empirically better understanding of substantive issues than OT, for example, of syllable typology in languages. Let us see how this works in the following section.

## 3 From Substance to Computation

### 3.1 The Case of Syllable Typology

#### 3.1.1 Syllable Typology in OT

We have argued that OT is problematic in terms of ‘the poverty of the inheritance’ in language evolution but that minimalist phonology is adequate both explanatorily and evolutionarily. So the required procedure for empirical verification is to show how achievements in OT are inherited to minimalist phonology and yet whether minimalist phonology is empirically more valid than OT. To achieve this goal, we take syllable typology as an example here.

Zec (2007: 165) offers a neat summary of the syllable typology of the world’s languages, as given in (2). An onset consonant is either required (R) or optional (O) while a coda consonant is either optional or banned (X), whose fact means that in general, the former is favored but the latter is disfavored.^4^ Consonant clusters in the onset and coda positions are also disfavored.

(2) Syllable Typology of the World’s Languages

| onset | coda | onset cluster | coda cluster | inventory | language | examples |
| --- | --- | --- | --- | --- | --- | --- |
| R | O | O | O | (C)CV(C)(C) | Totonak | CV, CCV, CVC, CCVC, CVCC, CCVCC |
|  |  |  | X | (C)CV(C) | Dakota | CV, CCV, CVC, CCVC |
|  |  | X | O | CV(C)(C) | Klamath | CV, CVC, CVCC |
|  |  |  | X | CV(C) | Temiar | CV, CVC |
| R | X | O | – | (C)CV | Arabela | CV, CCV |
|  |  | X | – | CV | Senufo | CV |
| O | O | O | O | (C)(C)V(C)(C) | English | CV, CCV, CVC, CCVC, CVCC, CCVCC, V, VC, VCC |
|  |  |  | X | (C)(C)V(C) | Spanish | CV, CCV, CVC, CCVC, V, VC |
|  |  | X | O | (C)V(C)(C) | Finnish | CV, CVC, CVCC, V, VC, VCC |
|  |  |  | X | (C)V(C) | Turkish | CV, CVC, V, VC |
| O | X | O | – | (C)(C)V | Pirahã | CV, CCV, V |
|  |  | X | – | (C)V | Fijian | CV, V |

R = required, O = optional, X = banned

In Totonak, for example, the onset is required, the coda is optional, and the clusters in the onset and coda positions are optional. So the resultant inventory is described as (C)CV(C)(C), including CV, CCV, CVC, CCVC, CVCC, and CCVCC.

Note here that there are sets of implicational relations in the inventory of syllable shapes of a language: the more marked (complex) shapes imply the more unmarked (simple) shapes, whose law is called “implicational universals” or “implicational markedness” (Jakobson 1958/1962, Clements and Keyser 1983, Prince and Smolensky 1993/2004). The most unmarked syllable shape is CV, and hence all the languages have CV other than more marked ones. (3) summarizes some implicational relations which are relevant here.

(3) Implicational Relations

a. Basic Typology Only CV. - - - Senufo CVC or V implies CV. - - - Pirahã, Fijian, Temiar, Turkish VC implies CVC and V. - - - English, Spanish, Finnish, Turkish
b. Complex Onset and Complex Nucleus CCV implies CV. - - - Arabela, Pirahã CVV implies CV. - - - languages with vowel length contrast CCV implies CVV. - - - languages with vowel length contrast
c. Complex Coda CVCC implies CVC. - - - Klamath, Finnish

Readers can examine these implicational relations by referring to the examples of the inventory in (2). In the syllable typology of (2), we do not take vowel length contrast into consideration, but it is attested that CVV implies CV and that CCV implies CVV. CVV depicts long vowels or diphthongs here, and the former implicational relation is obvious and seen in various languages (Blevins 1995). As for the latter, the complex onset is more marked than the complex nucleus, and all the languages with the complex onset in (2) also have long vowels and/or diphthongs. We will see how this is true in section 3.1.2.

Now let us see how these inventories and implicational relations are accounted for in OT, where language typology is captured as variation of ranking of given universal constraints. In this framework, constraints are violable, but violation must be minimal and only licensed under less dominant constraints to obtain optimality. For the basic typology in (3a), we assume that the input inventory is {CV, CVC, VC, V}. This analysis is based on the original account by Prince and Smolensky (1993/2004).^5^

(4) OT Analysis for Basic Typology

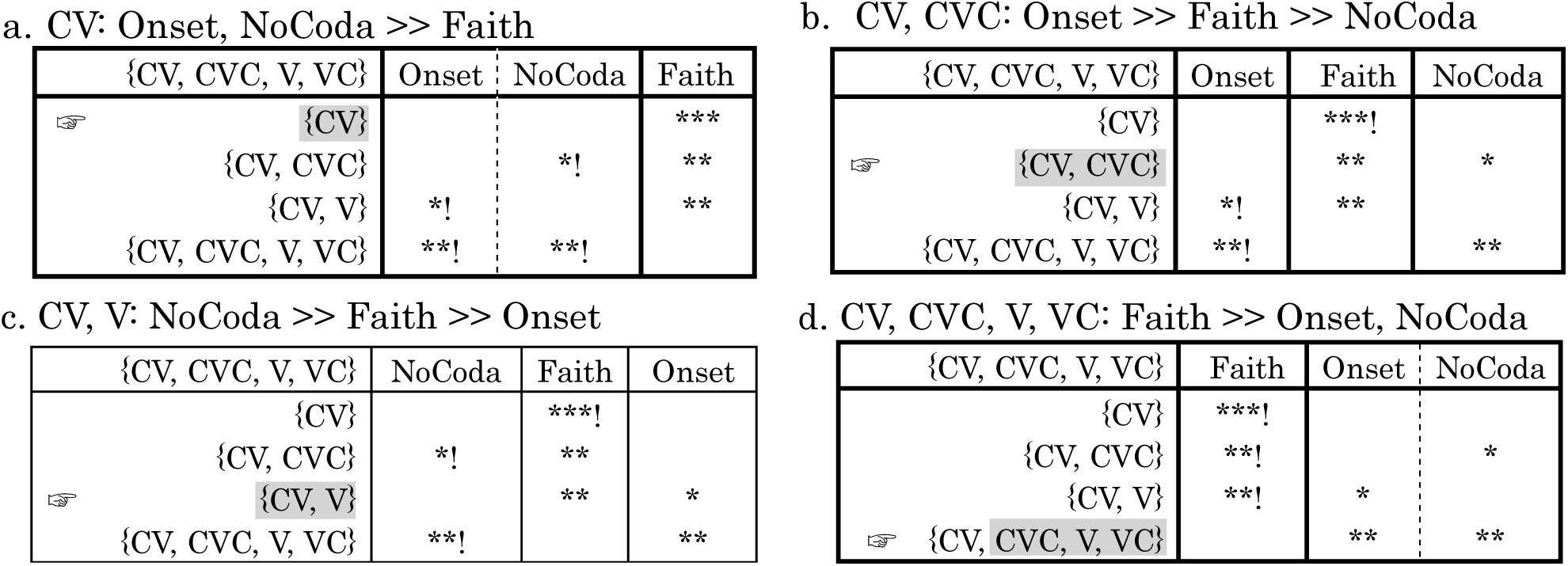

254 a. CV: Onset, NoCoda >> Faith
|  |  |  |  |  |
| --- | --- | --- | --- | --- |
| 255 | {CV, CVC, V, VC} | Onset | NoCoda | Faith |
| 256 | ☞ {CV} |  |  | *** |
| 257 | {CV, CVC} |  | *! | ** |
| 258 | {CV, V} | *! |  | ** |
| 259 | {CV, CVC, V, VC} | **! | **! |  |
| 260 c. CV, V: NoCoda >> Faith >> Onset |  |  |  |  |
| 261 | {CV, CVC, V, VC} | NoCoda | Faith | Onset |
| 262 | {CV} |  | ***! |  |
| 263 | {CV, CVC} | *! | ** |  |
| 264 | ☞ {CV, V} |  | ** | * |
| 265 | {CV, CVC, V, VC} | **! |  | ** |
| b. CV, CVC: Onset >> Faith >> NoCoda |  |  |  |  |
|  | {CV, CVC, V, VC} | Onset | Faith | NoCoda |
|  | {CV} |  | ***! |  |
| ☞ | {CV, CVC} |  | ** | * |
|  | {CV, V} | *! | ** |  |
|  | {CV, CVC, V, VC} | **! |  | ** |
| d. CV, CVC, V, VC: Faith >> Onset, NoCoda |  |  |  |  |
|  | {CV, CVC, V, VC} | Faith | Onset | NoCoda |
|  | {CV} | ***! |  |  |
|  | {CV, CVC} | **! |  | * |
|  | {CV, V} | **! | * |  |
| ☞ | {CV, CVC, V, VC} |  | ** | ** |

Here, an output inventory with any violation (*) of *dominant* constraints is crucially excluded (!) according to the ranking of the three constraints, and the resultant output candidate becomes an optimal inventory (☞). Since the three constraints allow only 6 possible permutations (i.e., 3!) and the reranking of Onset and NoCoda in (4a,d) does not change the results, the 4 patterns above are all and only inventories that the three constraints allow.^6^

In the same way, in the analysis for complex onset and complex nucleus, there are three constraints involved as in (5), and possible permutations are 6.^7^

(5) OT Analysis for Complex Onset and Complex Nucleus

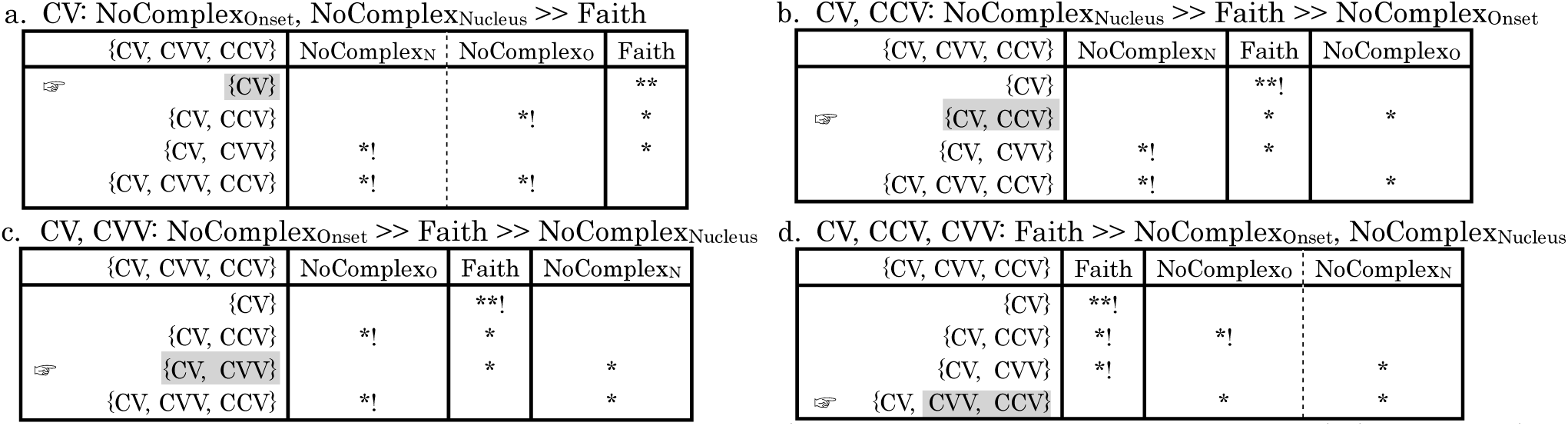

But again, the reranking of the two constraints NoComplex_Onset_ and NoComplex_Nucleus_ in (5a,d) is free and does not change the results, so the above 4 are acceptable as all and only relevant inventories. These 4 optimal inventories guarantee the implicational relations about complex onset and complex nucleus in (3b) as well as the basic shape without any complex onset or any complex nucleus. As for the complex coda case in (3c) where CVCC implies CVC, it can be accounted for by the ranking of Faith >> NoComplex_Coda_, which allows {CV, CVC, CVCC}. If the ranking is opposite and NoComplex_Coda_ is dominant, then the inventory of{CV, CVC} only becomes optimal.

#### 3.1.2 Syllable Typology in Minimalist Phonology

In minimalist phonology, there is no ranking of constraints or no evaluation on outputs by violable constraints. Instead, there are just interface conditions which are inviolable and apply after internalization (i.e., construction of hierarchical structure) and for externalization in the SM systems. Our account of syllable typology is based on the evolution and origins of language and hence draws an entirely different picture from OT.

Specifically, our account applies to the emergence of syllable shapes in phylogeny (biological emergence), ontogeny (developmental emergence), and glossogeny (historical or typological emergence). This is because the Third Factor as we discussed in (1) equally applies to the three aspects of evolution, which is repeated as (6).

(6) The Third Factor in Macroevolution and Microevolution (= (1))

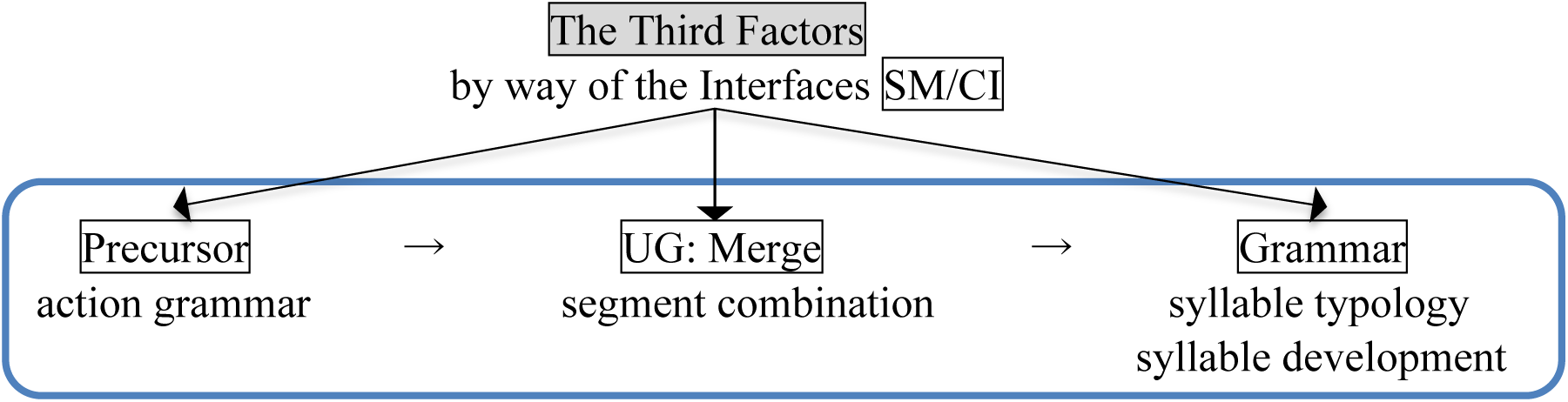

A syllable is a combination of segments (vowels and consonants) and has an internal organization that is formed by Merge. On one hand, the development and typology of syllable shapes in grammars stem from the on/off settings of the interface conditions on Merge-based combinations in the SM systems for externalization (section 3.3). On the other hand, Merge stems from some precursor: we assume here following Fujita (2016a, 2017) that Merge has been preadapted or exapted from action grammar in sense of Greenfield (1991). This idea is called the Hypothesis on Motor Control Origin of Merge in Language Evolution.

Although Merge seems uniquely human, its emergence is not a miracle or skyhook but gradual according to this hypothesis. Considering a developmental and evolutionary connection between action and language, Greenfield distinguishes three methods of object combination, as in the case of cup nesting given in (7a-c), and focuses especially on the crucial step from the pot-type (7b) to the subassembly-type (7c) of manipulations of objects. Analogous structures in language are given on the right, in which X means the host or head of that combination as compared to a larger cup.

(7) Action Grammar and Language

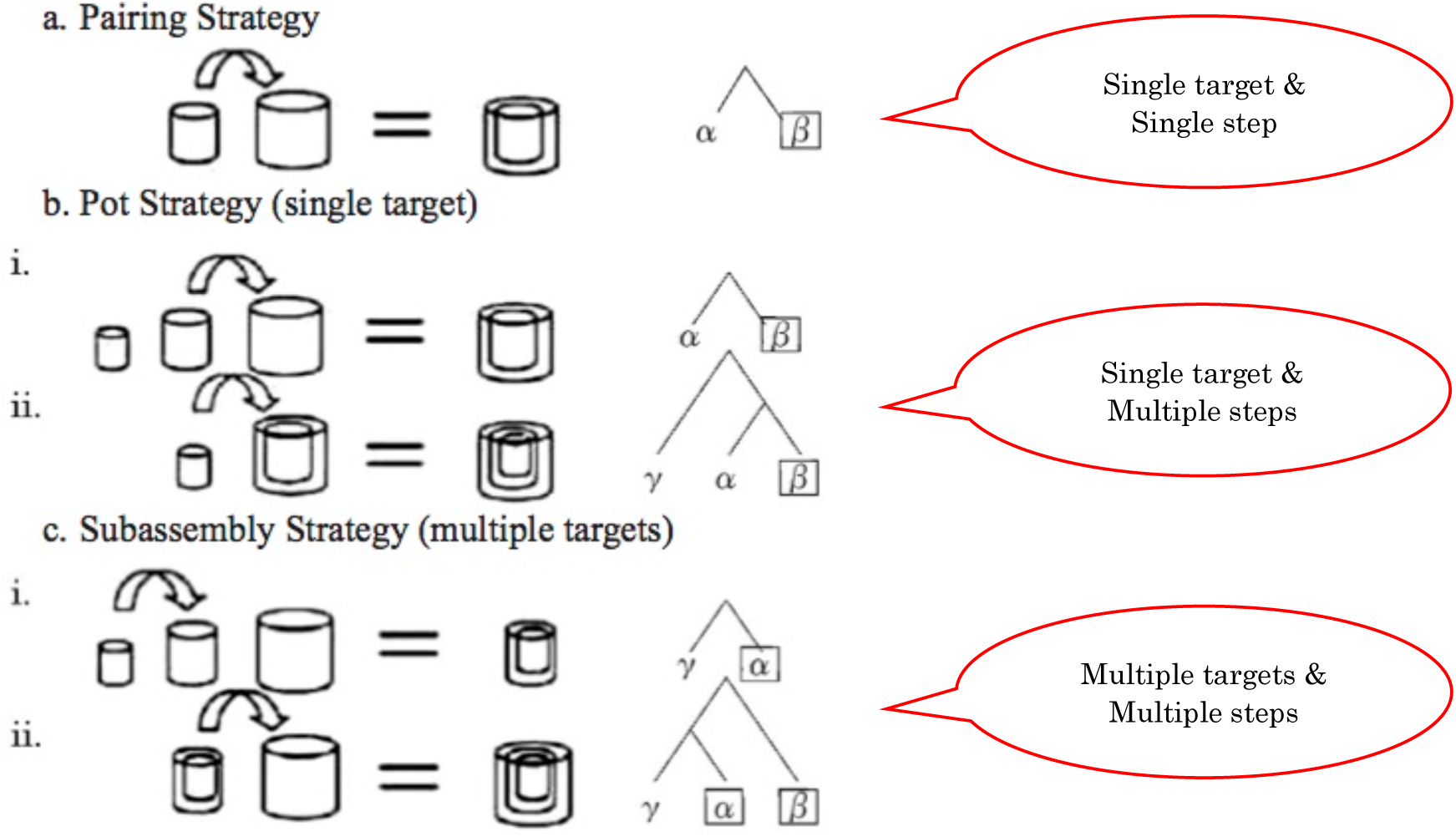

Both the pot and subassembly strategies may be said to be recursive operations in that the combined objects as a result of one nesting 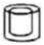 are carried on to become the input to the next nesting with another object. Then, both strategies can create hierarchical structures in the case of language, and the final results are the same 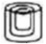 in the case of cup nesting. So it is the process of each operation that is crucial here, not the result of each process.

In the pot strategy there is only a single target (host or head) that is combined with other objects, whereas in the subassembly strategy there are two targets (hosts or heads): a previous subunit (= subassembly) with one target is combined with another. In that sense, the subassembly strategy requires more complex chunking and more working memory than the pot strategy (see Aboitiz et al. 2006 for the connection between combination type and working memory). In fact, it is reported from a comparative cognitive experiment by Maynard Smith and Szathmáry (1995) that only the pot strategy is observed in chimpanzees while the subassembly strategy is used by human infants as young as 20 months old. So only the subassembly strategy is uniquely human. Then in Fujita’s (2016a, 2017) scenario of parallel evolution between action and language, the uniquely-human subassembly strategy in action was somehow transferred by preadaptation or exaptation into sub-Merge in language, which is exactly the origins of human language in macroevolution.^8^ In this way, the species-specific aspect of Merge has not emerged out of nothing but gradually.

Note that the Third Factor, the implicational law (the more complex implies the simpler) here, applies to action and language alike: the subassembly strategy is implied by the pot strategy, which is in turn implied by the pairing strategy. Now let us see how the inventories and implicational relations in (3) are accounted for in our view. First, the basic typology in (3a) is derived as in (8). All languages must take the first step of pair-Merge and that is why they have CV consistently.^9^

(8) Merging Processes in Basic Typology

a. pair-Merge: first step

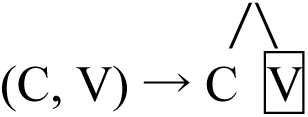
b. unMerge: V implies CV.

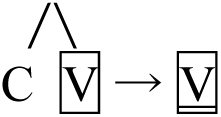
c. pot-Merge: CVC implies CV.

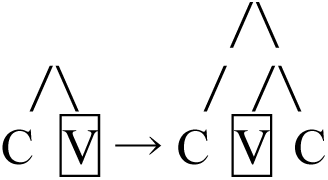
d. pot-Merge followed by unMerge: VC implies CVC (and CV).

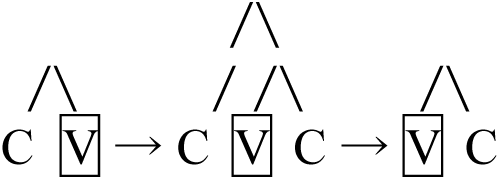

UnMerge followed by pair-Merge: VC implies V (and CV).

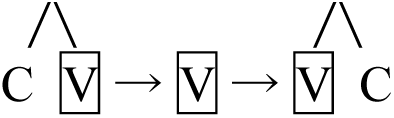

UnMerge in (8b,d) is the reverse or cancelling process of a previous operation of Merge, or pruning of the non-head element of a Merged pair. It remains left for future research whether it is a cancelling process of Merge or a pruning process applied by some interface condition. Here we assume that it is cancelation of a previous application of Merge and that it requires the same effort and working memory as pot-Merge, because the head element concerned is only one in both operations.

As for merging processes for complex onset, nucleus, and coda, the derivation for (3b,c) is accounted for by the following way; the emergence of complex onset and complex coda involves sub-Merge while that of complex nucleus only requires pot-Merge.

(9) Merging Processes in Complex Onset, Nucleus, and Coda

a. sub-Merge: CCV implies CV.

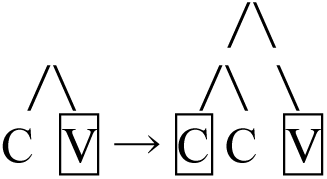
b. pair-Merge followed by pot-Merge: CVV implies CV.

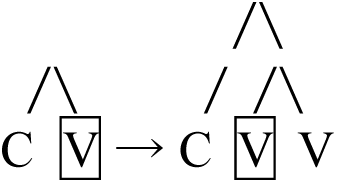
c. pot-Merge followed by sub-Merge: CVCC implies CVC (and CV).

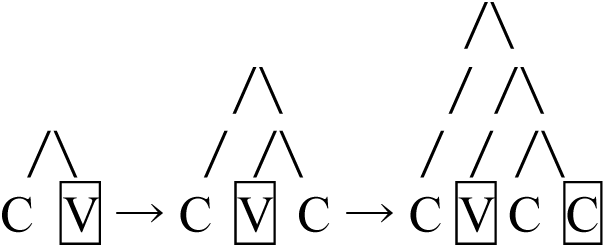

The case that remains to be explained in (3b) is the one in which CCV implies CVV. This fact simply follows from the implicational law by which the more complex implies the simpler. Since the derivation of complex onset requires sub-Merge as in (9a) and that of complex nucleus requires only pot-Merge as in (9b), it is quite natural that the former implies the latter.

In this way, the implicational law as the Third Factor works on glossogeny, the historical change of language, as well as on phylogeny and ontogeny and forms the typology of syllable shapes in the world’s languages. However, there might be some apparent exceptions. That is, it is not always true that a sub-Merged structure implies a pot-Merged structure in the same operation of a language, *under a certain condition*. For example, Fujita (2016b) discusses an interesting case: Japanese has both direct passives (derived by sub-Merge) and indirect passives (derived by pot-Merge), while English only has direct passives.

But this is due to the fact that direct passives are evidently superior to indirect ones in processing active/passive alternation and that indirect passives has lost or replaced with direct ones. In general, a pot-Merged structure may be replaced with the emerging sub-Merged structure in glossogeny *if the latter has a superior function in usage to the former*. In contrast, if there is no superiority/inferiority in function between the two, then they can coexist. For example, we have seen that CVCC implies both CVC and CVVC, but English has CVC (e.g., *hit* via pot-Merge) and CVVC (e.g., *heat* via pot-Merge) as well as CVCC (e.g., *hint* via sub-Merge). This is because the three syllable have no superiority/inferiority to each other in meaning contrast. So when we look into implicational relations in language, it is to be noted that covert (implicit) implication does not always become overt (explicit) in the real aspects of typology.

### 3.2 Crucial Empirical Difference: Superheavy Syllables

Now let us examine a more complicated case of superheavy syllables CVCC and CVVC, which involve complex coda and complex nucleus, respectively. These two actually stand in an irreversible implicational relation: languages with CVCC also have CVVC but not vice versa (Tanaka 2018b, Fujita, Taniguchi, and Tanaka to appear).

(10) Implicational Markedness for Superheavy Syllables

a. CVCC and CVVC English, Totonac (MacKay 1991), Klamath (Barker 1963, 1964), Finnish (Keyser and Kiparsky 1984), and so on.
b. Only CVVC Japanese, Karukv (Bright 1957), Yup’ik (Jacobsen 1984), Leti (Engelenhoven 1995), and so on. Thus, there are no languages with CVCC and without CVVC in syllable typology, and the fact that languages with CVCC also have CVVC means that the quantity contrast in the coda implies the one in the nucleus.

In investigating the implicational relation between CVCC and CVVC, it is important to examine whether there is independent evidence showing that CVCC or CVVC in question is ‘truly’ superheavy. For example, English has a quantity contrast in final stress between heavy and superheavy syllables, as shown in (11) (Hayes 1980/1981, Halle and Vergnaud 1987).

(11) English: Final Stress on CVCC and CVVC

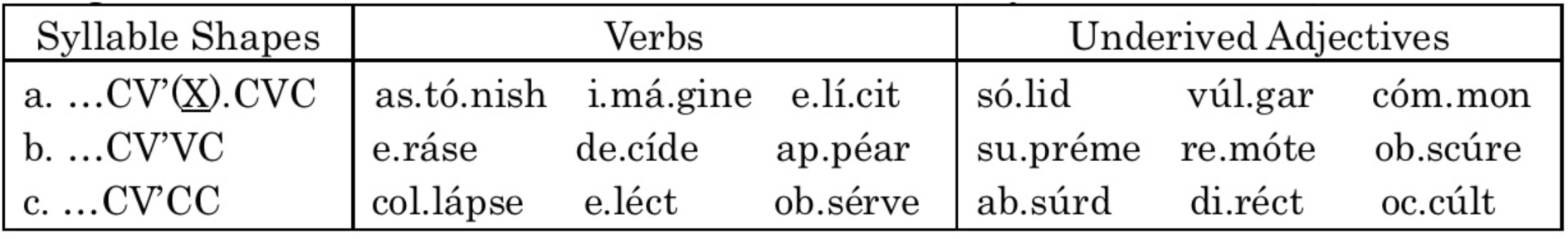

CVCC and CVVC in (11b,c) attract final stress while CVC in (11a) does not, so they are not simply heavy but must be superheavy.^10^ On the other hand, in Japanese, there is no CVCC because this language does not allow complex coda, but there is evidence for the status of CVVC as a superheavy syllable.

(12) Japanese: High-Level Tone on CVVC

| Syllable Shapes | Adverbs and Nouns |  |
| --- | --- | --- |
| a. <u>CV</u>CV... | zu.rut.to ‘slipped’ | zi.rit.to ‘impatiently’ |
|  | ko.mu.gi.ba.ta.ke ‘wheat field’ | mu.zin.sai.to ‘unattended site’ |
| b. CVV. ... | zuu.to.pi.a ‘Zootopia’ | zii.mee.ru ‘G-mail’ |
|  | koo.mu.in ‘civil servant’ | muu.tai.ri.ku ‘Mu Continent’ |
| c. CVVC. ... | zuut.to ‘for a long time’ | ziin.to ‘sympathetically’ |
|  | koon.ba.ta.ke ‘corn field’ | muun.rai.to ‘moonlight’ |

If CVVC in (12c) were not superheavy but split into two consecutive syllables CV.CV, then it would exhibit low-rising tone just like (12a). Instead, CVVC forms a syllable domain and exhibits high-level tone in the same way as CVV in (12b) (Tanaka 2000, 2013). It then follows that CVVC is not a sequence of CVs but a superheavy syllable.

Given the fact that CVCC implies CVVC in syllable typology, our account is straightforward from the viewpoint of minimalist phonology. As we have discussed so far, implicational relations follow from 1) the derivational steps of Merge and 2) the complexity of the mode of Merge in syllable construction. In particular, languages using sub-Merge for syllable construction use pot-Merge as well, but not vice versa: languages using pot-Merge may not use sub-Merge. This implicational law is reflected on the difference in internal organization between CVCC and CVVC.

Superheavy Syllables: CVXC

a. pot-Merge followed by sub-Merge

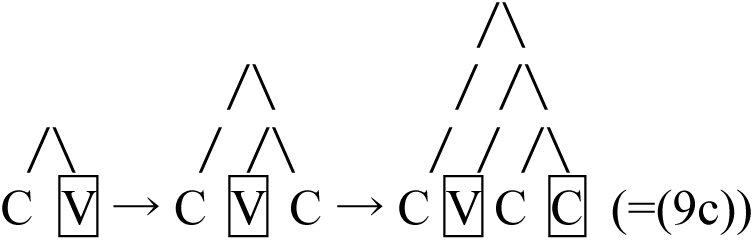
b. pot-Merge followed by another pot-Merge

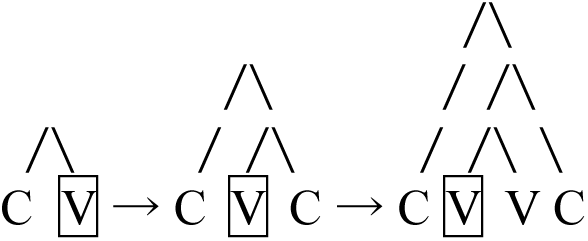

As shown in (13), the derivation of CVVC requires pot-Merge, but that of CVCC requires sub-Merge. This is why CVCC implies CVVC and not vice versa. Recall that CCV implies CVV as was shown in (3b), whose relation is accounted for in the OT tableau of (5d). In our account, the derivation of CCV requires sub-Merge as in (9a) and that of CVV requires only pot-Merge as in (9b). and so the former implies the latter.

As for complex margins, it is generally true that “[i]f more than one consonant is allowed in a margin, there is in principle no limit to the number permitted” (aside from phonotactics) (Zec 2007: 164). In fact, it often happens that languages with complex onset or complex coda have three- or four-consonant cluster. This is a direct consequence of the fact that sub-Merge is required to derive the complex onset for CCV as in (9a) and the complex coda for CVCC as in (13a). However, it does not hold true that if more than one vowel is allowed in a nucleus, there is no limit to the number of vowels permitted. According to Blevins (1995: 217), languages with three-way contrast for vowels (i.e., CV vs. CVV vs. CVVV) are quite rare. This is because the derivation of vowels in a nucleus requires just pot-Merge whose operation is limited and does not involve recursion. In this sense, recursion should be characterized by sub-Merge, and recursive Merge is nothing but sub-Merge.

Unfortunately, however, factorial typology in OT predicts that there would be languages with CVCC and without CVVC, as is evident in (14c).

(14) OT Analysis for Superheavy Syllables

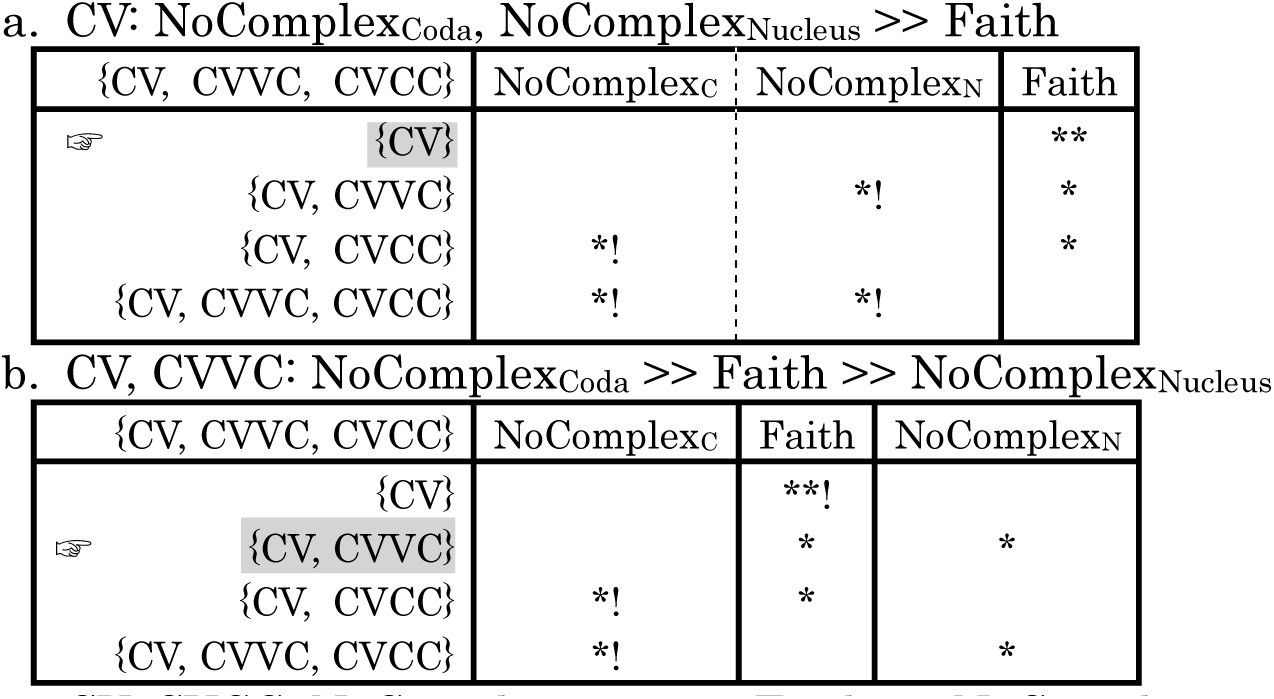

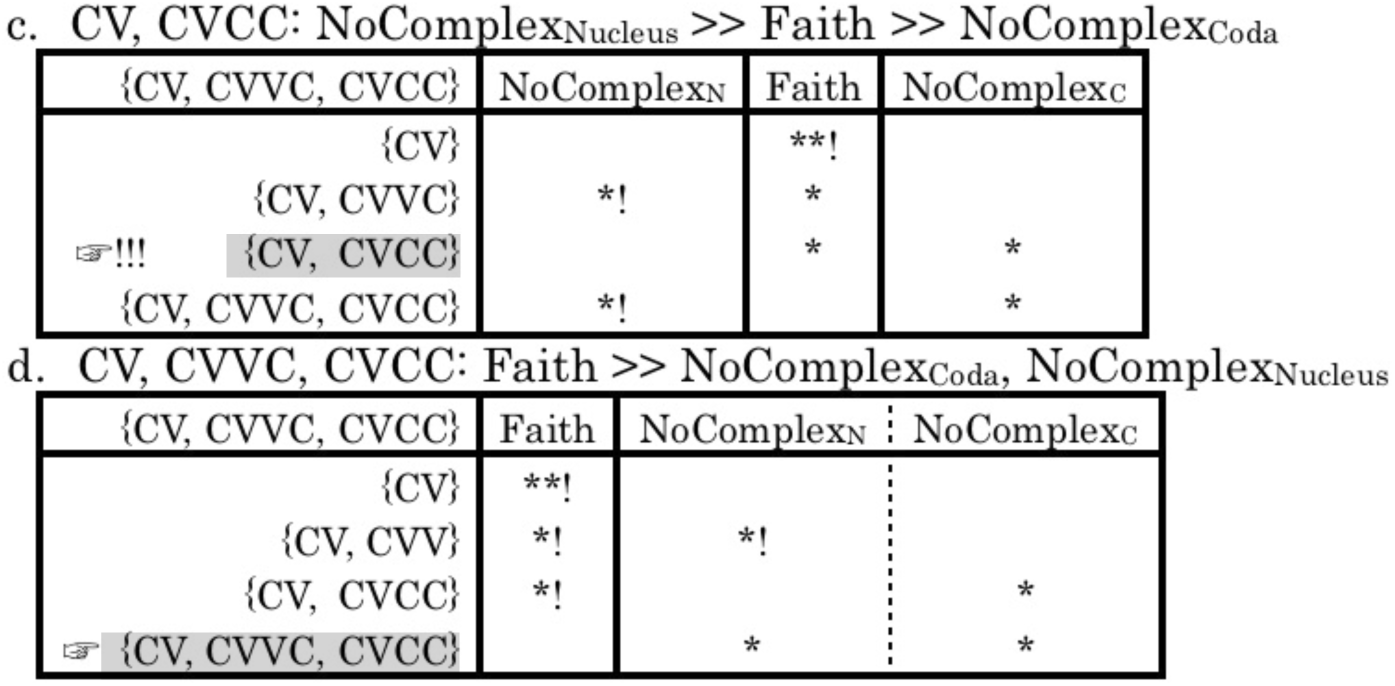

Then as far as syllable typology is concerned, minimalist phonology is empirically more valid than OT while the achievements in OT can basically be inherited to minimalist phonology. For arguments against OT from other points of view, see Hale and Reiss (2008: 191-254).

### 3.3 The Place of Constraints and Substantive Issues in Minimalist Phonology

We now have seen from the discussion so far that there are no constraints in UG since it involves only Merge. So there are no CON (constraint set and ranking), GEN, and EVAL in UG like OT. Yet constraints are still working at the SM interfaces in externalization or linearization. The so-called ‘constraints’ in OT are ‘interface conditions’ in minimalist phonology, and substance is all about issues of the SM systems at the interface, which are subject to the Third Factor.^11^ The Third Factor involves principles of the physical and natural law, such as implication, efficiency, and economy, which work for the mappings from segment combinations via Merge to the grammar of syllable structure.

(15) The Role of Interface Conditions

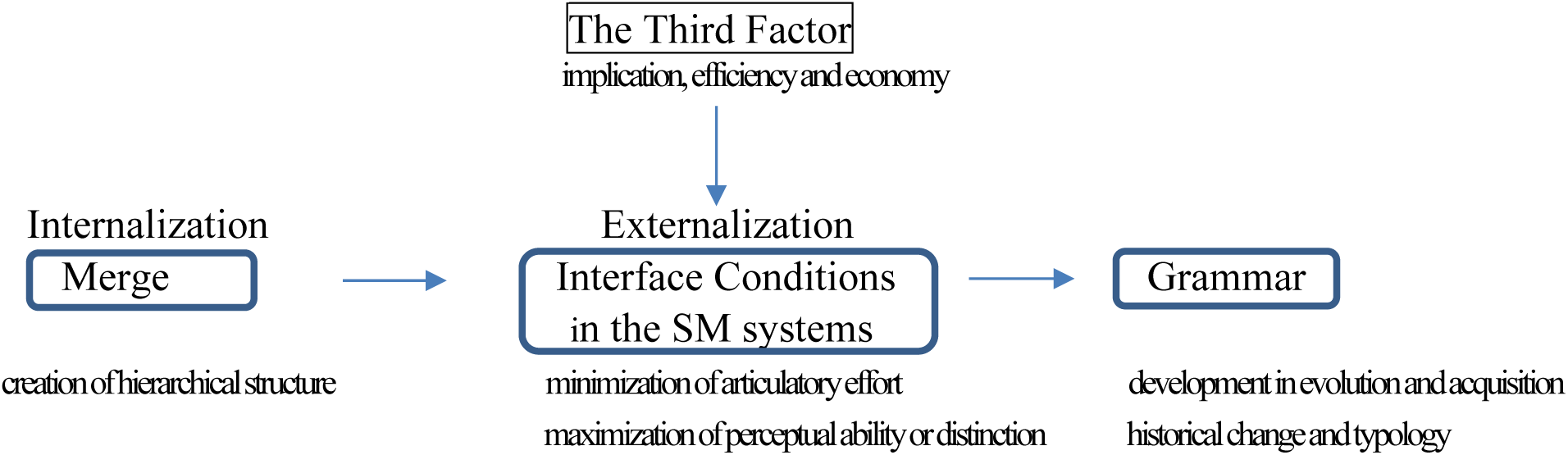

Given this architecture of grammar, then, how are substantive issues such as acquisition, social variation, and historical change accounted for as well as typology? This is the question we now have to answer.

Take the origin of syllables, i.e., the phylogeny of syllable structure, as an example. As we stated in (8a), the initial state of syllable phylogeny is CV. This fact seems to indicate that the settings of interface conditions, including Onset and NoCoda, are [on] at this stage. The same is true for the initial state of syllable ontogeny, and the acquisition of syllable structure begins with CV, changing in shape from the simpler to the more complex (Levelt, Schiller, and Levelt 2000, Levelt and Vijver 2004). That is why the implicational law works. So syllable construction begins with CV, not V, VC, or VV as in (16a), because all interface conditions are [on] in the initial state of ontogeny (language acquisition) and phylogeny (language evolution).

(16) The Shift in Settings of Interface Conditions

a. pair-Merge

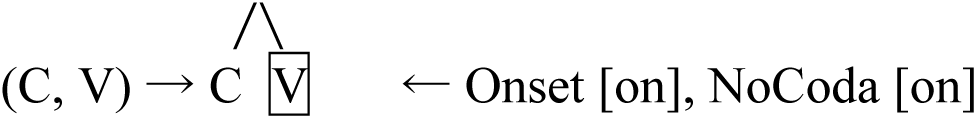
b. unMerge

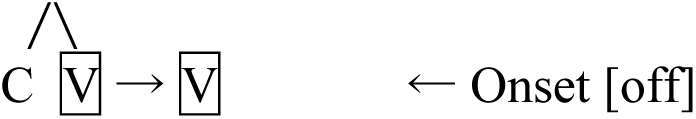
c. pot-Merge

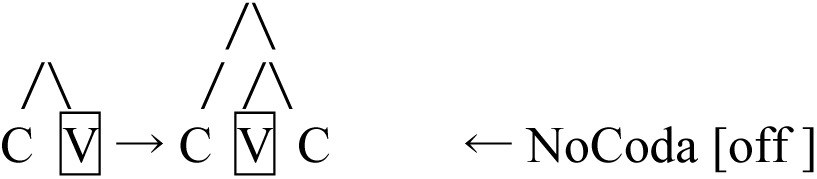
d. pot-Merge followed by unMerge

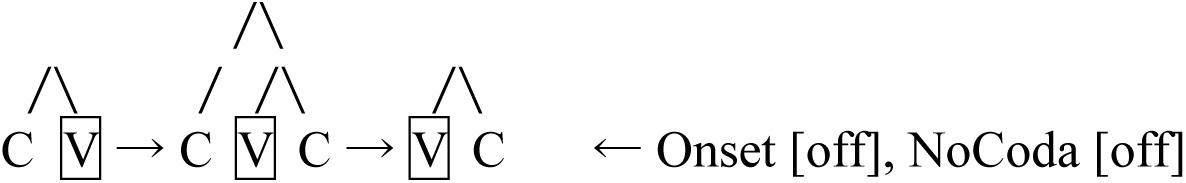

UnMerge followed by pair-Merge

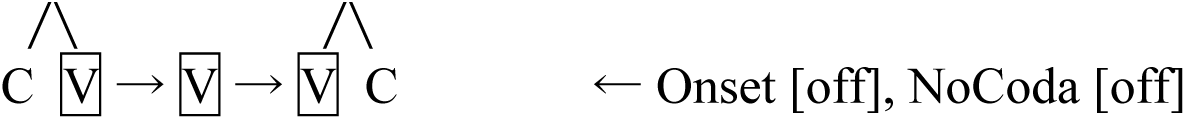

In minimalist phonology, OT constraints are redefined as interface conditions on a merged pair of phonological units in externalization/linearization. That is, all conditions are stated *in terms of the linear order of phonological units*. In the case of syllables, the relevant conditions are redefined as in the following way.

(17) Interface Conditions on Syllable Shapes

a. Onset: in a merged pair (C, V), C must be followed by V.
b. NoCoda: in a merged pair (C, V), V must not be followed by C.
c. NoComplex: in a merged pair (C, C) or (V, V), C or V must not be followed by the same element.

Then, language typology follows from possible shifts of the initial [on] to [off] of the interface conditions, as in (16b), which lead to language variation and change. Possible shifts are due to environmental and social factors, and they might work as ‘parameters.’ The state of affairs in syllable typology can schematically be shown with the settings of the interface conditions as in (18).

(18) Settings of the Interface Conditions in Syllable typology.

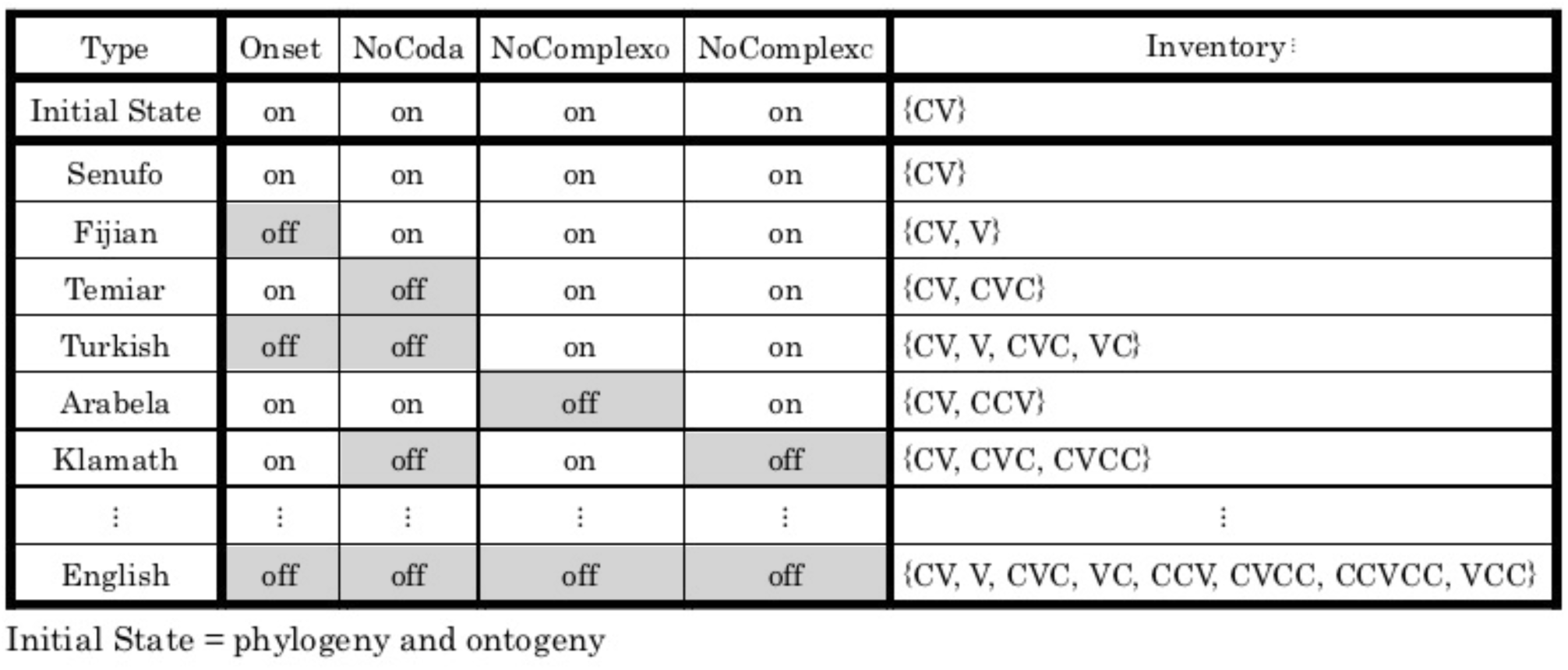

Recall that in (2), the possible syllable shapes to be covered are 12 types. Given the [on/off] settings of the four conditions in (18), possibilities are calculated as 2×2×2×2, resulting in 16 types.^12^ However, NoCoda [on] necessarily implies NoComplex_Coda_ [on], as a coda cluster is absolutely prohibited if there should be no coda consonant, and hence there are 4 impossible types (2×2) with NoCoda [on] and NoComplex_Coda_ [off]. As a consequence, possible syllable shapes in (18) amount to (2×2×2×2)-4 = 12 types, which agree with Zec’s (2007) syllable typology.

In summary, substantive issues including typology, acquisition, historical change, and social variation are accounted for by the [on/off] settings of interface conditions in (19), which were issues over (re)ranking of markedness and faithfulness constraints (abbreviated here as M and F) in OT.

(19) Substantive Issues in Minimalist Phonology

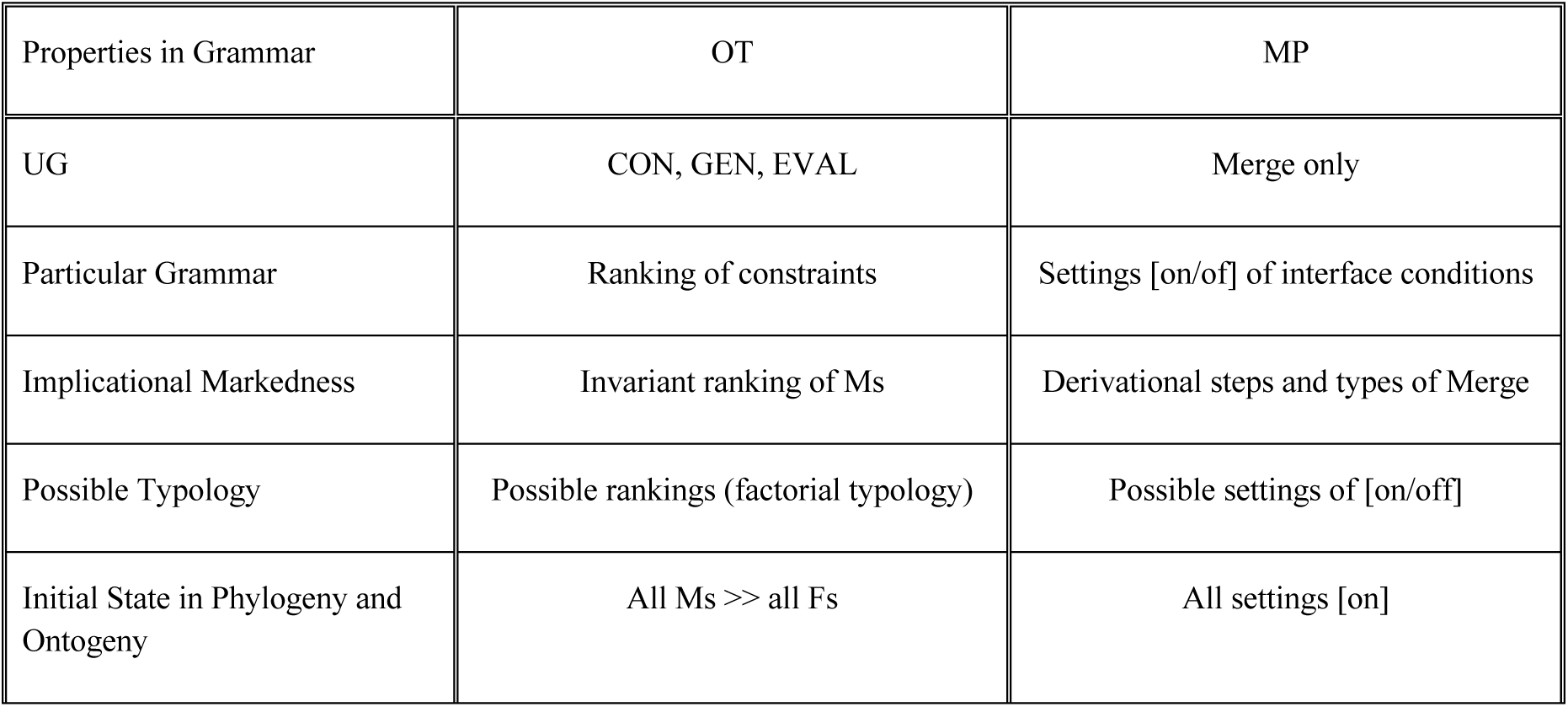

|  |  |  |
| --- | --- | --- |
| Acquisition in Ontogeny (to the marked) | Demotion of Ms | Shift from [on] to [off] |
| Ultimate Historical Change (to the Marked) in Glossogeny | Demotion of Ms | Shift from [on] to [off] |
| Proximate Historical Change (to the Unmarked) in Glossogeny | Demotion of Fs | Shift from [off] to [on] |
| Synchronic Variation | Free ranking of M and F | Free setting of [off] or [on] |

Note here that there are two aspects in glossogeny, which are not to be confused. One is a macro-level, ultimate, historical change that has evolved from the initial state of phylogeny, and the other is a micro-level, proximate, historical change that has developed from the initial state of ontogeny. In our view, the former ultimate change is the shift from the simpler to the more complex in which some interface conditions are set to [off], bringing about language variety and typology in a very long time frame. The latter proximate change is the shift from the complex to the simpler in which some interface conditions are set to [on], bringing about simplification in pronunciation, or minimization of articulatory effort from one generation to the next. It is during this proximate change in progress that the synchronic variation of a single form occurs, when both the [off] and [on] settings coexist in the older and younger generations.

In this way, OT’s findings about substantive issues can successfully be carried over to minimalist phonology. Yet minimalist phonology is more valid than OT from the viewpoint of ‘the poverty paradox’ and the typology of syllable shapes.

## 4 Ambiguity: Evidence for Internalization and Hierarchical Structure in Phonology

In the previous sections, we have shown that the internal organization of a syllable is based on the three types of Merge and that this view offers an empirically better account of syllable typology than OT. The internal organization of a syllable is a typical hierarchical structure, just like that of a sentence. In general, binarity, headship, and dominance count as characteristics of the hierarchical structure of human language, and they may cause structural ambiguity. So structural ambiguity is often said to show evidence for the hierarchical structure, and disambiguation of the same linear string requires internalization, precisely because externalization alone cannot make a distinction among the linearly ambiguous structures. In this way, internalization plays a crucial role in computation, a core part of grammar, and is nothing but a species-specific and domain-specific property in human language, unlike the substantive issues of externalization in the SM systems.

Then, if phonological structure is hierarchical, it may well exhibit structural ambiguity. This means that phonology involves internalization and that it is not just a function for externalization. On the other hand, it is generally believed that, as Chomsky (2008: 136) notes, phonology is “an ancillary process … doing the best it can to satisfy the problem it faces” at the interface of the SM systems, meaning that it is just an issue in externalization. Even on the phonology side, Samuels (2011: 34, 59) states that “the operations and representations underlying phonology were exapted, or recruited from other cognitive domains for the purpose of externalizing language” and that “the human phonological system is a domain-general solution to a domain-specific problem, namely the externalization of language.” In this view, no part of phonology is unique to language or humans, or for that matter, is part of internalization, a virtually consistent claim with Chomsky.

However, it is evident that syllable structure is hierarchical, exactly because a wide array of implicational relations follows from its internal organizations and offers a valid account of syllable typology. Furthermore, syllable structure exhibits structural ambiguity, which is observed in the phonological string CCVC. Let us see how ambiguity and disambiguation work on this structure.

It is generally true that in English, the same consonant appears on the <u>C</u>V<u>C</u> string as in *pop, Bob, tit, dad, kick, gag, sis, rare, nun, mom, lol*, and so on, while it cannot appear on the C<u>C</u>V<u>C</u> string and there are no such words as *\*plol, *prare, *blol, *brare, *trare, *drare, *twew, *dwew, *clol, *crare, *glol, *grar, *snun, *smom, *swaw, *slol, *srare, *flol, *frare*, and so on. This is because the C<u>C</u>V<u>C</u> string has a peculiar identity avoidance effect between the second onset and the coda, forming a systematic gap (Baertsch 2002, Baertsch and Davis 2003).^13^ However, a syllable with the so-called sC cluster (the cluster of /s/ and a voiceless stop) is definitely free from this effect, since there are such actual words with C<u>C</u>V<u>C</u> as *stat* and *stet*, which forms a crucial contrast with the above mentioned *\*snun, *smom, *swaw, *slol*, and *\*srare*.

Now *stat* and *\*snun* have another difference: the sC cluster violates the Sonority Sequencing Principle, but the other clusters of /s/ and a nasals/liquid/glide conform to it, forming a valid syllable. So concerning the sC cluster, the aberrant segment /s/ lies outside the domain of a valid syllable and is adjoined directly to the syllable node, not belonging to the onset node.

(20) Identity Avoidance Effect on *[[CC_i_][VC_i_]], not on [s[C_i_[VC_i_]]]

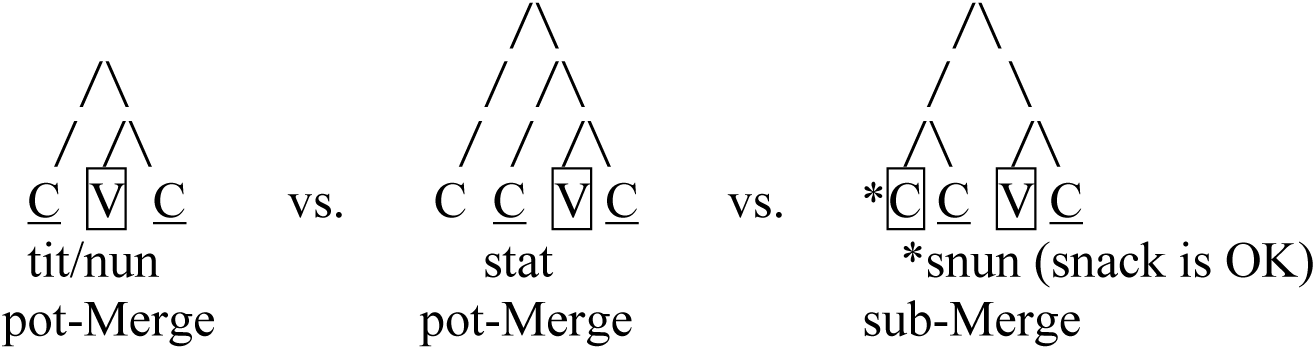

This internal disambiguation among the two types of CCVC that have the same linear order has further evidence from the ontogeny of the English syllable. At the initial stage of syllable acquisition, it is very difficult for infants to produce consonant clusters, and they have different strategies for resolution depending on the cluster types: the sC cluster is simplified by deleting /s/ as in *(s)tay, (s)tory, (s)poon*, and *(s)pend*, while valid onset clusters including the /s/-initial ones undergo deletion of the inner sonorants as in *s(n)ack, s(m)all, s(l)eep, s(w)im*, and the like (Smith 1973, 2010). In short, cluster simplification reflects the difference in internal organization between the two types of clusters and applies by deleting the more sonorous of the two consonants in each type.^14^

This disambiguation seems difficult to state in terms of interface conditions in the SM systems alone; for example, a statement fails like “linearization clashes if a coda consonant is preceded by the same second onset within the domain of CCVC.” So the identity avoidance effect cannot be defined *only* in linear segmentation, but the hierarchical structure is crucial. Phonology is not exclusively a matter of the SM systems in externalization/linearization but may involve some mechanism in internalization. In this way, syllable structure offers evidence for the hierarchical structure below the word level.

## 5 Conclusions

In summary, the role of phonology in the SMT is to probe into the following two issues: the substantive issue of how phonological phenomena are accounted for by Merge (computation) and interface conditions (substance), and the computational issue of how phonology can offer evidence for the hierarchical structure in human language through Merge.

The title “From substance to computation” means “from OT (substance-centric) to minimalist phonology (computation-centric).” In the history of phonological theory, it is computation that has been crucial for a better understanding of substance and for each stage of its paradigm shift. In this sense, computation is the core issue in phonological theory. In minimalist phonology, computation lies in the core component of Merge, and substance is drawn from the SM systems at the interface. Importantly, minimalist phonology can resolve the apparent paradox between the richness of UG in acquisition and the poverty of UG in evolution, which means that it has explanatory and evolutionary adequacy.

It is also true for OT that computation is crucial. For OT to be a theory of computation in grammar and undergo due paradigm shift, phonologists should focus on the architecture of GEN and EVAL in UG, including issues like the controversy between parallelism and serialism. However, OT is problematic in terms of evolutionary adequacy, because UG has a rich content of CON, GEN, and EVAL. Also, it suffers from an empirical problem with syllable typology. These two problems are ultimate and proximate issues in substance. Since the adequacy of computation is empirically tested by ultimate and proximate issues in substance, OT needs to reexamine its mode of computation.

Instead, computation in minimalist phonology can inherit the achievements of OT, and yet the former is empirically more valid than the latter, for example, in the case of syllable typology. The [on/off] settings of the interface conditions can account for substantive issues not only in typology but also in language acquisition or language variation and change.

So further questions that remain to be answered are i) how various substantive issues are accounted for in minimalist phonology, ii) if phonology is not a matter of the SM interfaces alone, what evidence there is for internalization in phonology, and iii) if it is a theory of computation, how the whole hierarchical structure is created by Merge with the interface conditions.

## 6 Conflict of Interest Statement

The author declares that the research was conducted in the absence of any commercial or financial relationships that could be construed as a potential conflict of interest.

## 7 Author Contributions

The author confirms being the sole contributor of this work and has approved it for publication.

## 8 Funding

This research is supported by a MEXT Grant-in-Aid for Scientific Research on Innovative Areas #4903 (Studies of Language Evolution for Co-creative Human Communication) whose specific branch is “Theoretical Frameworks for Studying the Origins and Evolution of Language” (subject number: 17H06379) and by a Grant-in-Aid for Basic Science (B) of the Japan Society for the Promotion of Science whose specific research topic is “Internal and External Evidence for Phonological Theory: A Transdisciplinary Approach” (subject number: 16H03427).

## 9 Acknowledgments

Part of the contents of this article was publicized at an invited talk entitled “Rethinking syllable typology from the perspective of evolinguistics: From universal constraints to interface conditions” in the Tokyo Conference on Evolinguistics 2018: “Where Does Phonology Fit In?” held on March 7, 2018, at the University of Tokyo, Japan. I would like to express my deepest gratitude for valuable comments and feedback from the audience, especially to Pedro Martins, Kuniya Nasukawa, and Bridget Samuels. Any errors left are the sole responsibility of the author.

## Footnotes

1 Here, “ultimate issues” denote biological ones in macroevolution, and “proximate issues” historical or typological ones in microevolution.

2 There have actually been a few occasions where its computational systems are reexamined. One is the proposal of the shift from Containment Theory to Correspondence Theory in which evaluation is based not only on outputs but also on input-output or output-output relations (McCarthy and Prince 1995). Here we will argue against Correspondence Theory as a standard version of OT. Another is Harmonic Serialism (McCarthy 2008, McCarthy and Pater 2016), which frames a hybrid computational system in which the ‘serial’ mode of derivation is revived and incorporated into OT’s ‘parallel’ evaluation. For arguments against Harmonic Serialism, see Fujita, Taniguchi, and Tanaka (to appear).

3 We should say in strict terms that “the uniquely-human trait is only *a certain form of Merge*,” since primitive forms of Merge may be shared with other species. This point will be discussed in section 3.2.

4 There is no language whose onset is banned systematically, unlike the coda or the clusters.

5 Here, Onset is a constraint that requires a consonant in the onset position, and NoCoda is a constraint that prohibits a consonant in the coda position. Faith is a correspondence constraint that charges as many violation marks as any difference between the input and output inventories.

6 Even if the input inventory were changed from{CV, CVC, VC, V}, the results would be the same in the four patterns and guaranteed by the ranking of the constraints.

7 NoComplex is a constraint that prohibits complex structure in the margins and the nucleus. Its family includes NoComplex_Onset_, NoComplex_Coda_, and NoComplex_Nuclues_, depending on the relevant position.

8 Hereafter, the pairing, pot, and subassembly strategies in action are called pair-Merge, pot-Merge, and sub-Merge, respectively, in language. See Boeckx and Fujita (2014) for a debate over the parallel evolution between action and language, and for some arguments against skeptic views.

9 As we will discuss in section 3.3, all the interface conditions are [on] in the initial state of phylogeny and ontogeny. So even if strings such as CC, VV, and VC emerge at the first step, derivation clashes due to conditions like Onset, NoCoda, and NoComplex. This is why CV is the only choice.

10 It happens to nouns that CVV attracts final stress as in *employée* and *ballét*, which does not mean that CVV is superheavy. Instead, this is just a case of morphologically-driven final stress and limited to some peculiar suffixes such as *-ee* and *-et*.

11 What we call ‘interface conditions’ include markedness constraints in OT, which require minimalization of articulatory effort in motor control. There are no conditions that are equivalent to faithfulness constraints, but some interface conditions require maximization of perceptual ability or distinction in the sensory systems.

12 We do not take the quantity of vowels, i.e., NoComplex_Nucleus_, into consideration, just like the typology in (2).

13 This identity effect is not observed on the (s)<u>C</u>CV<u>C</u> string as in *plip, blob, twit, dread, click, quick, grog, strut, street*, and *straight*, or on the (s)<u>C</u>VC<u>C</u> string as in *pump, bulb, tilt, damned, kirk, spump, start, stunt, skunk*, and *skulk*.

14 The fact that for a valid syllable, the target of deletion is the more sonorous consonant of the onset cluster shows that the onset position favors the less sonorous one and that the outer consonant is the head of the onset in *snack, small, track, plot, crush*, and so on. That is why the outer onset C of the third syllable structure is boxed in (20). The difference in internal organization between the two types of clusters in (20) is further supported by the observation in Smith (2010: 214-215) that tri-consonantal clusters such as *straight, strawberry*, and *strong* develop in the way of *(s)t(r)*V → *(s)tr*V → *str*V. In general, deletion of the non-head in syllable acquisition means that the development of syllable structure in ontogeny is characterized as attachment of the emerging non-head to the existing head position, and this may also be true for its development in glossogeny and phylogeny. This picture conforms to our idea developed in section 3.1.2.

## Notes

### Competing Interest Statement

The authors have declared no competing interest.

